# Ontogeny of catechol-o-methyltransferase expression in the rat prefrontal cortex: effects of methamphetamine exposure

**DOI:** 10.1101/2025.02.06.636913

**Authors:** Lauren K. Carrica, Joshua M. Gulley

## Abstract

Repeated use of methamphetamine (METH) is known to dysregulate the dopaminergic system and induce long-lasting changes in behavior, which may be influenced by sex and age of exposure. Catechol-o-methyltransferase (COMT) is an enzyme that is involved in the breakdown of catecholamines, and its role in dopamine clearance is thought to be especially important in the prefrontal cortex (PFC) where dopamine transporter (DAT) expression is relatively scarce. The first study in this report utilized a rat model to characterize the ontogeny of COMT protein expression in the PFC and nucleus accumbens (NAc) across adolescence, which is a developmental stage that has been shown to involve significant reorganization of dopaminergic innervation. Drug-naïve male and female Sprague-Dawley rats were sacrificed on postnatal day (P) 29, 39, 49 or 69, and expression levels of COMT protein within the PFC and NAc were analyzed via Western blot. We found that COMT expression in the PFC increases across adolescence in a sex-dependent manner but does not significantly change in the NAc during this timeframe. A separate group of rats were injected daily from P40-48 (adolescence) or P70-78 (adulthood) with saline or 3.0 mg/kg METH and sacrificed on P49 or P79. While METH decreased COMT in adult rats of both sexes, METH increased COMT expression in the PFC of rats exposed in adolescence. The results of this work suggest that exposure to METH during adolescence uniquely effects dopamine clearance within the PFC, potentially contributing to differences in neurobiological outcomes from METH use.

## Introduction

Adolescence, which in humans occurs from approximately 12-20 years of age, is a period of significant physical, behavioral, and cognitive changes. For example, the prefrontal cortex (PFC) undergoes substantial reorganization from adolescence into young adulthood [1][2][3][4] that is characterized by considerable changes in dopamine signaling. Axons immunoreactive for tyrosine hydroxylase, the rate-limiting enzyme in the synthesis of dopamine, increase innervation into the PFC during adolescence in rats [5] and in nonhuman primates [6][7]. While the total volume of the PFC declines in both humans [1][8] and rodents [9], dopamine inputs to the PFC, as measured by tyrosine hydroxylase immunoreactivity, increase in a sex-specific way during adolescence to levels that are significantly higher than those seen either earlier or later in life [6][10]. Regulated dopaminergic signaling within the brain is critical for numerous cognitive and motor functions and disruption within these pathways has been demonstrated to play a role in a number of neurological diseases, including substance use disorders (SUDs) [11].

Methamphetamine (METH) acutely increases dopamine levels within various regions of the brain [12], and changes to dopamine signaling represent one of the critical components of the behavioral effects of METH and its potential to lead to compulsive use and relapse that contributes to SUDs [13]. While SUDs are problematic at any life stage, stimulant misuse may be of particular concern when it impacts the developing brain. Over 60% of people who use stimulants other than cocaine report initiating use prior to age 18, and adolescent-onset users are 70% more likely to develop the clinical symptoms of SUD within 2 years compared to adult-onset users [14]. In addition, adolescent METH users are far more likely than their non-using peers to exhibit risky sexual behavior, polydrug use, and other behavioral problems [15][16]. The exact neurobiological underpinnings of these differences are difficult to establish in humans, due in part to the myriad of factors that may contribute to substance misuse during adolescence.

However, a growing body of work in animal models suggests that differences in key aspects of dopamine signaling play a pivotal role in determining outcomes from drug use. The dopamine system is a leading candidate mechanism for the unique effects of adolescent compared to adult exposure to amphetamines because of the delayed development in the PFC and the potent effect amphetamines have on dopaminergic signaling within this region [17][18].

While the changes in dopamine innervation and signaling has been investigated in the context of adolescence, what is less often studied is the developmental trajectory of dopamine clearance mechanisms. The primary mechanism of dopamine clearance from the synapse is via the dopamine transporter (DAT), which returns dopamine into the presynaptic cell where it can be metabolized by monoamine oxidase (MAO) or repackaged into synaptic vesicles [19]. In the PFC, however, DAT expression is much lower than in other areas of the brain [20]. Due to this, the enzyme catechol-o-methyltransferase (COMT) plays a particularly crucial role within the PFC. COMT is produced in two different isoforms: the short, soluble form (S-COMT) that is thought to be involved mostly when catecholamine levels are high [21][22][23] and the longer, membrane-bound form (MB-COMT) that is primarily involved in the termination of dopaminergic signaling when catecholamine levels are lower and more physiologically relevant [24]. One study in rats demonstrated that COMT is responsible for approximately 60% of dopamine clearance within the PFC [25]. This critical role of COMT may be enhanced in the instance of METH use, as one of the effects of METH is the inhibition [26] and reversal [27][28] of the DAT such that it releases dopamine into the synapse via a mechanism referred to as exchange diffusion.

In humans, there are three functional polymorphisms of COMT that differ based on the Valine (Val) or Methionine (Met) allele at codon 158 of the COMT promoter region: Val/Val, Val/Met, and Met/Met. The valine allele is hypothesized to confer relatively higher COMT activity compared to the methionine allele, such that the Val/Val variant would be expected to confer the highest COMT activity and Met/Met the lowest [29]. Polymorphisms of the COMT gene have been associated with altered cognition assessed in the PFC-sensitive Wisconsin Card Sorting Test [30]. Additionally, certain polymorphisms of this gene have been linked to increased incidence of SUD diagnosis for several drugs [31], including alcohol [32] and METH [33][34].

The effects of COMT expression on SUD may be age- and gender/sex-specific. In both humans (gender) and mice (sex), extreme COMT reduction (via knockout in mice and through 22q11.2 deletion syndrome in humans) is associated with cortical thinning only in females and only after puberty. This same study demonstrated that both humans and mice showed COMT-by-sex effects in executive function that were paralleled by alterations to tyrosine hydroxylase and that only emerged in adolescence [35]. These sex and age effects may furthermore play a role in substance use and outcomes following cessation. In humans, female Val/Val carriers report more severe urges to smoke and withdrawal symptoms following overnight abstinence from nicotine than Val/Met carriers. This effect is not seen in men [36]. In rats, the COMT inhibitor tolcapone reduces both reinforcer seeking and alcohol consumption in male, but not female, alcohol-preferring “P” rats [37]. Taken together, these results suggest that the effects of COMT activity may be influenced by both age and gender/sex, and that these factors may all interact to lead to diverging outcomes of drug seeking and withdrawal effects. It is currently unknown, however, how COMT expression or function is changing during the same period of adolescent development when other aspects of the dopamine system are still maturing.

To establish whether COMT expression is changing in sex-specific ways across adolescence, we used a rat model of adolescence to investigate the ontogeny of COMT protein expression in the PFC and nucleus accumbens (NAc) from early adolescence through early adulthood. Time points were selected such that measures before and after puberty in both male and female rats were taken since pubertal onset has been demonstrated to be a critical point of transition in neurodevelopment [38]. Once the general pattern of COMT expression was determined, we investigated the effects of repeated exposure to METH during adolescence or young adulthood to determine if drug-induced changes in COMT expression were dependent on age of exposure. We hypothesized that COMT would follow a similar developmental trajectory to tyrosine hydroxylase [5] in that expression would peak in the PFC following pubertal onset in both males and females and plateau into young adulthood. Because of its relatively reduced functional significance in the earlier-developing NAc, we predicted there would be no age-related changes in this brain region. Furthermore, we expected that METH exposure would increase expression of both isoforms of COMT in the PFC in all groups in response to repeated elevations in dopamine levels both intra- and extra-cellularly, but that the increase would be greater in adolescent-exposed rats.

## Methods

### Subjects

A total of 96 Sprague-Dawley male and female rats were used for these experiments (n = 24/sex for the ontogeny experiment and n = 24/sex for the drug exposure experiment). Rats were housed in same-sex groups of three beginning on postnatal day (P) 23, were kept on a 12:12 light/dark cycle (lights off at 0900), and were provided food and water *ad libitum.* Rats were checked daily beginning on P29 for physical markers that indicate pubertal onset: preputial separation for males and vaginal opening for females [39][40]. In the sample of rats used here, mean pubertal onset was 35.3 ± 0.8 days for females (range: 32-38 days) and 43.2 ± 0.4 days for males (range: 41-45 days). All animals were handled and weighed daily in the hour before dark cycle onset (∼0800). In both the ontogeny and the drug exposure experiments, rats were assigned randomly to groups for the most equal distribution of littermates across treatment and/or age groups.

### COMT ontogeny: Tissue collection and estrus cycle determination

For the drug-naïve rats used to determine the ontogeny of COMT expression (n = 24/sex), sacrifice occurred at one of four ages (n=6/age/sex): P29, 39, 49, or 69. All post-pubertal females in the ontogeny groups underwent a vaginal lavage after anesthesia and immediately before decapitation to assess estrus cycle stage. Stage of estrus cycle (proestrus, estrus, or diestrus) was determined using Goldman’s methods as previously described [41]. Animals were anesthetized with 2.5% isoflurane for 2-3 min before sacrifice via rapid decapitation. Following decapitation, brains were extracted in ≤ 5 min, immediately frozen in dry ice, and subsequently stored at −80 °C until bilateral micropunches were taken from 350-μm coronal slices of the medial PFC (prelimbic and infralimbic regions combined; bregma point AP +3.72 to +2.0) and the nucleus accumbens (bregma point AP +2.28 to +1.2). Punches were taken only where anatomical markers confirming proper placement (e.g. anterior forceps of the corpus callosum for the PFC and lateral ventricles/anterior commissure for NAc) were visually confirmed to reduce the chances of differential sampling across ages.

### Effects of METH on COMT expression

Rats used in this experiment (n=24/sex) were injected (i.p.) with 0.9% saline or 3.0 mg/kg METH (Research Triangle Institute; distributed by The National Institute on Drug Abuse) in a volume of 1 ml/kg/injection. These injections occurred daily within the first hour of the dark cycle (∼0900-1000h) for 9 consecutive days, either from P40-P48 (adolescent exposed) or P70-78 (adult exposed). The side of the body where injections occurred was alternated daily to avoid discomfort around injection sites. Rats were sacrificed 24-26 hours after their final injection (P49 or 79) using the procedure described above. Brain extraction and tissue processing also occurred as described above, except only samples of the PFC were taken.

### Western Blot Analysis

Samples were homogenized and lysed in icy cold lysis buffer (150mM NaCl, 50mM Tris-44 HCL pH = 7.4, 30mM EDTA, 1.5% Triton-X, 0.1% SDS, 0.5% protease inhibitor and 0.5% phosphatase inhibitor) using a standardized protocol previously described [42]. Protein concentration was estimated for each sample using Precision Red Advanced Protein Assay (Cytoskeleton, CA). Samples were prepared at 20 μg protein/well with 4X laemmli loading buffer and 2-mercaptoethanol, heated at 95C for 5 min, and run electrophoretically on pre-cast gels (4-20% TGX Stain-Free Protein Gels, Bio-Rad) with at two samples from each same-sex group on each gel. Samples from male and female animals were run on separate gels. Following electrophoresis, protein was transferred to PVDF membrane (Bio-Rad), and blocked in 5% non-fat milk in TBST overnight at 4. Strips of membrane were incubated in 5% non-fat milk in TBST for 1.5 hours at room temperature in primary antibody [COMT: 1:2500, ProteinTech] and secondary antibody for 2 h at room temperature. Vinculin was used as the housekeeping protein (1:5000, ProteinTech). Images were captured using a ChemiDoc system (Bio-Rad) and densitometric analysis was performed using ImageJ (NIH, Bethesda, MD).

### Data Analysis

For each gel, the mean gray value (MGV) from COMT bands was normalized to the MGV value of the vinculin bands. An average of the same-sex adult control group within each gel was then calculated as the baseline to account for the variability across gels within this group. Since data from each sex were normalized separately to their respective control group, male and female data were analyzed separately, and thus sex is not a factor included in our ANOVA. In addition, COMT isoforms were analyzed separately rather than as a within-subjects factor because the goal of this study was to present the developmental trajectory for both S- and MB-COMT as independent variables. For the ontogeny experiment, one-way ANOVA (age) was used to analyze group differences in COMT expression in both the PFC and NAc. In the METH exposure experiment, two-way ANOVA (age x treatment) was used to analyze group differences in COMT expression within the PFC. All analyses were conducted in R (Version 1.4.1717; rstudio.com). Significant main effects were further analyzed via Tukey’s HSD.

## Results

### COMT ontogeny

To establish the ontogeny of COMT across adolescence, drug-naïve animals were sacrificed at one of four time points spanning early adolescence into young adulthood, and COMT protein expression was measured in both the PFC and NAc. One-way ANOVA revealed that in both males and females, expression of MB-COMT significantly increased between P29 and P39 in the PFC (male: F_3,20_ = 10.4, *p* < 0.001; female: F_3,20_ = 14.8, *p* < 0.001). In males (Fig.1), levels of MB-COMT did not significantly change from P39 through adulthood. In contrast, females (Fig. 2) showed a significant decrease in MB-COMT expression from P49 to P69 (F_3,20_ = 24.8, *p* < 0.001). In female animals, there was no significant difference between P29 and P69 (F_3,20_ = 1.4, *p* = 0.271). For animals of both sexes, MB-COMT expression within the NAc did not change significantly across age.

**Figure 1.**
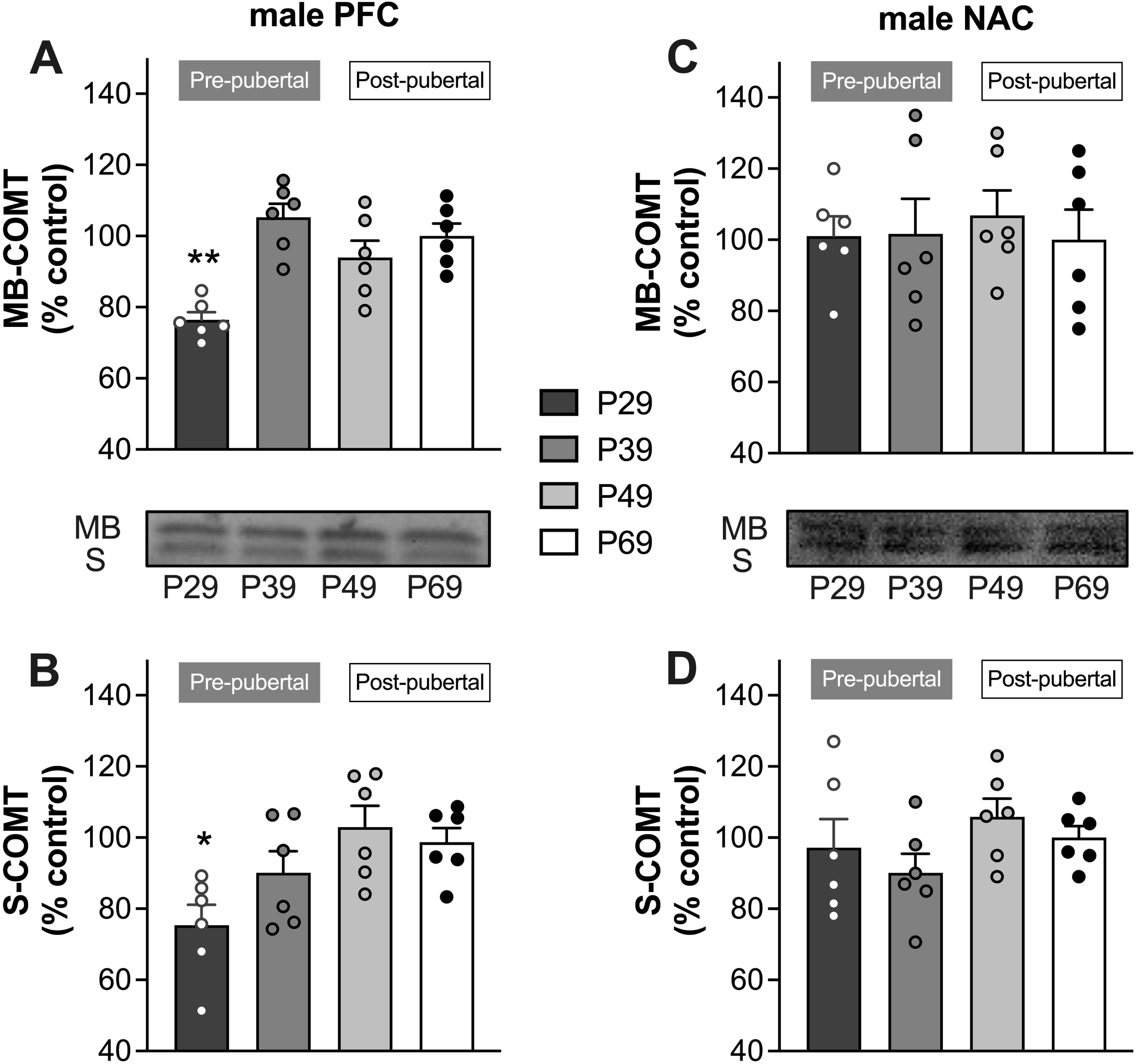
COMT expression across adolescent development in the PFC (A-B) and NAc (C-D) of male rats. Data are expressed as a percent of protein expression in adult males (control) for membrane bound (MB)- and soluble (S)-COMT, which are labeled on the representative Western blot image. Brain tissue was collected from different rats (n = 6/age group) that were pre-(P29 and P39) and post-pubertal (P49 and P69). \**p* < 0.05 vs. adult*, **p* < 0.01 vs. adult

**Figure 2.**
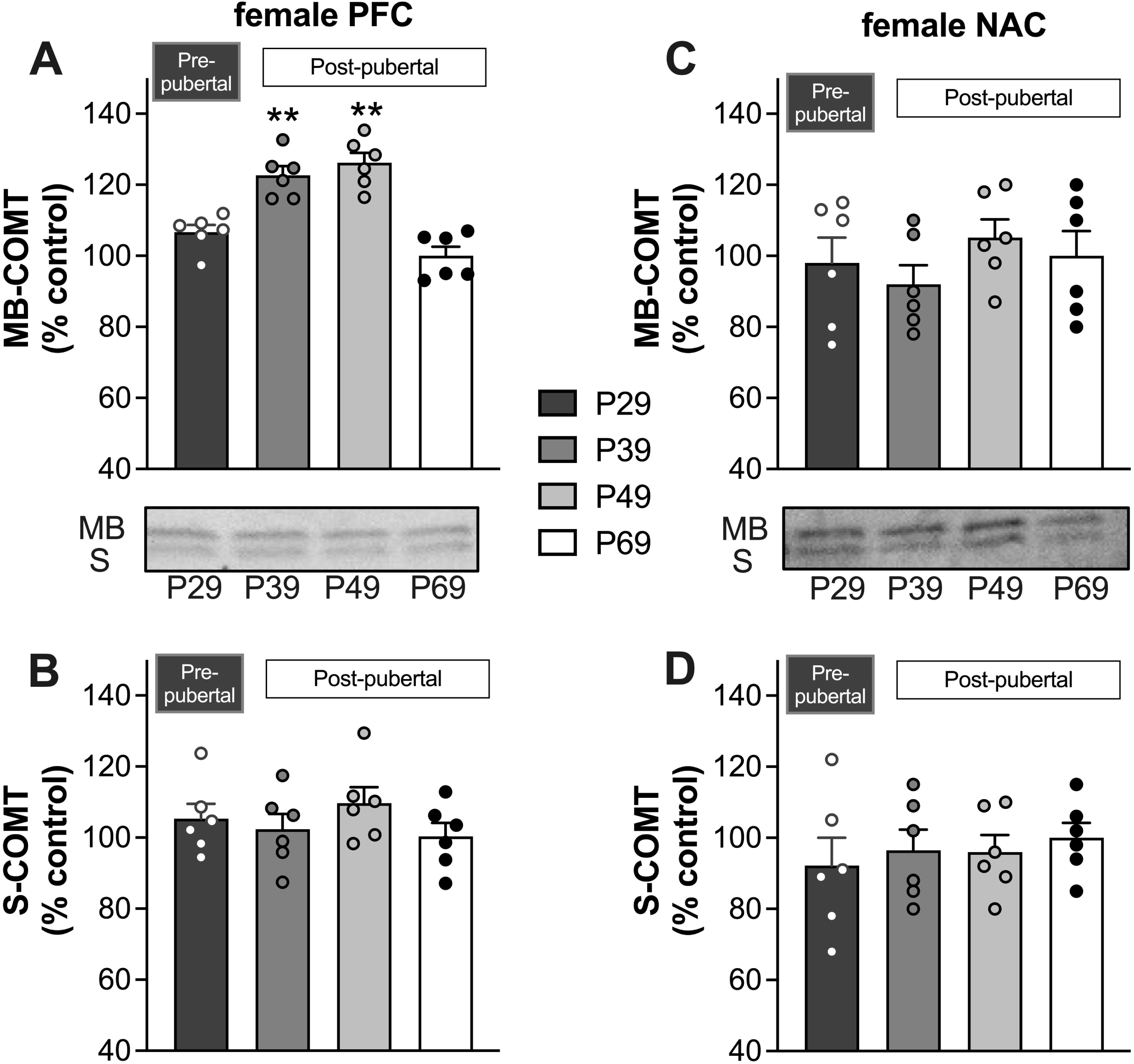
COMT expression across adolescent development in the PFC (A-B) and NAc (C-D) of female rats. expressed as a percent of protein expression adult females (control). Data are expressed as a percent of protein expression in adult males (control) for membrane bound (MB)- and soluble (S)-COMT, which are labeled on the representative Western blot image. Brain tissue was collected from different rats (n = 6/age group) that were pre-(P29) and post-pubertal (P39, P49, and P69). *\*\*p* < 0.01 vs. adult

For S-COMT, only male animals showed a significant change in expression across age (Fig. 1). In males, the change in S-COMT within the PFC expression mirrored that of MB-COMT in that expression increased between P29 and P39 (F_3,20_ = 4.86, *p* = 0.01) and remained consistent through P69. In females (Fig. 2), no significant changes in S-COMT expression in the PFC were detected (F_3,20_ = 0.938, *p* = 0.441). In the NAc, S-COMT did not change significantly across age in either male or female rats.

### Effects of METH on COMT expression

To investigate whether ongoing development of COMT expression during adolescence leads to unique effects of drug exposure on COMT expression, we injected adolescent (P40) or adult (P70) rats with saline or 3.0 mg/kg METH once daily for 9 days. We found that in both sexes, the effect of METH on COMT expression in the PFC was age specific. In female rats (Fig. 3A,B), METH significantly increased expression of both MB- and S-COMT in adolescent-exposed animals (F_1, 20_ = 12.1, *p =* 0.008 and F_1, 20_ = 12.6, *p =* 0.002, respectively), but in adult-exposed rats it reduced expression of both COMT isoforms (F_1, 20_ = 7.10, *p* = 0.034 and F_1, 20_ = 6.32, *p =* 0.04). Likewise in males (Fig. 3C, D), METH significantly increased expression of both MB- and S-isoforms (F_1, 20_ = 5.98, *p =* 0.04 and F_1, 20_ = 6.18, *p* = 0.042, respectively) in adolescent-exposed rats, but lowered expression in those exposed in adulthood (F_1, 20_ = 6.88, *p =* 0.039 and F_1, 20_ = 7.08, *p =* 0.03. In females, there was also a significant age difference in COMT expression in saline-injected controls, with expression of both isoforms significantly higher at P79 compared to P49 (F_1, 20_ = 6.36, *p =* 0.042 and F_1, 20_ = 6.17, *p =* 0.046, for MB- and S-COMT respectively). This age-dependent difference in controls was not observed in males.

**Figure 3.**
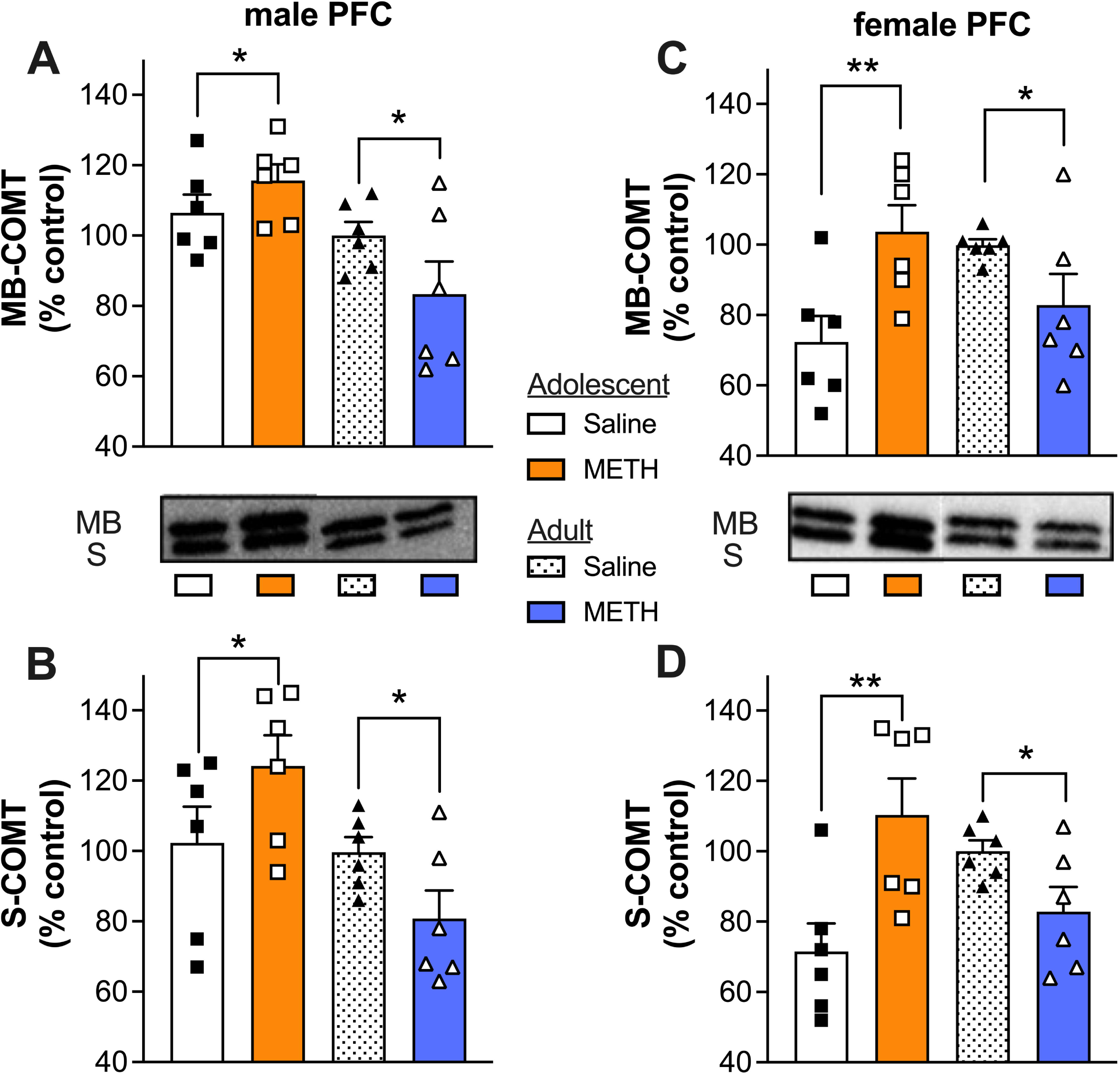
Effects of METH on MB- and S-COMT expression in the PFC of males (A-B) and females (C-D). Data (n = 6/group) are shown as a percent of protein expression in the adult, same-sex controls (saline exposed). \**p* < 0.05 and \*\**p* < 0.01 vs saline-injected controls ^#^*p* <0.05 vs adult saline.

## Discussion

Consistent with our hypothesis, COMT expression levels within the PFC changed across development in both male and female rats, indicating that dopamine clearance mechanisms beyond just the DAT undergo significant changes across adolescence just as dopamine innervation does [5][10]. In male animals, there was an increase in both isoforms of COMT at P39, relative to P29 that remained elevated through early adulthood (P69). In females, the change in COMT expression was isoform specific. MB-COMT significantly increased from P29 to P39 and remained elevated at P49, followed by a return to juvenile-like expression levels at P69. S-COMT did not significantly change in female animals at any time point measured.

Unlike the changes in the PFC, we did not find a significant change in COMT expression across any of the tested ages in the NAc. Dopaminergic neurons in the mesolimbic pathway project into the NAc, and dopamine signaling into this region is critical for the reinforcing properties of drugs like METH [43][44]. In contrast to the PFC, however, the DAT is highly expressed within the NAc and DAT is the primary mechanism of dopamine clearance within this region [45]. Furthermore, COMT expression does not appear to be as tightly linked to extracellular dopamine levels in the NAc as it is in the PFC. One study that examined extracellular dopamine levels in COMT knockout mice found a significant increase in dopamine within the PFC, but not in dorsal striatum or the NAc of either male or female animals [25].

The meaning of the differences in the developmental patterns of COMT isoforms is less clear. These two isoforms are thought to have at least somewhat distinct roles within the body. MB-COMT may be primarily involved in the termination of dopaminergic neurotransmission when there are low, physiologically relevant concentrations of catecholamines [24]. This isoform has also been shown to have a higher affinity for catechol substrates – and for dopamine in particular – than S-COMT. S-COMT, in contrast, is thought of as the high-capacity isoform [46]. S-COMT may therefore play an important role in the elimination of excessive, and potentially toxic, catecholamine levels that are induced by exogenous stimuli like METH. One limitation in studying COMT expression is that there are not yet antibodies that reliably distinguish S- and MB-COMT isoforms, limiting the ability to say whether their distribution across cell types or layers within the PFC is different. Separation by weight through Western blot allows for analysis of these two isoforms, but additional work is needed to determine the underlying differences in their maturational courses.

To determine if ongoing development of COMT signaling within the PFC may render adolescent animals particularly vulnerable to METH-induced changes in COMT expression, we next exposed animals of both sexes to METH or saline in either adolescence or adulthood. We utilized a dosing paradigm that we have previously demonstrated induces opposing effects on parvalbumin (PV) number in adolescent and adult females [47], an effect that may be influenced by differences in dopaminergic signaling given that PV+ interneurons rely on proper dopaminergic input to develop their normal adult function [48]. We began dosing for adult animals at P70 when rats would presumably have baseline COMT expression levels comparable to what we demonstrated at P69 rather than beginning at P60 when the difference seen between P49 and P69 females may not be complete. Animals were sacrificed ∼24 h following their final drug exposure, allowing us to investigate short-term changes to COMT signaling after blood levels of METH have decreased significantly. We hypothesized that METH exposure would significantly increase COMT expression in both age groups, but to the greatest extent in adolescents. Interestingly, we found that METH exposure significantly increased COMT expression within the PFC in adolescence, but decreased expression of COMT in adulthood. This was true for both isoforms of COMT and for both sexes, unlike our previously reported effects on PV which were specific to females. These results suggest that COMT may contribute, at least in part, to sex differences in METH-driven changes in PV expression between adolescents and adults. There appears to be a sex-specific factor that makes females uniquely vulnerable, at least at this age and dosing regimen.

There has been a significant amount of literature describing the effects of altering COMT activity on the effects of amphetamines [49][50], alcohol [51][52], and nicotine [53][54], but very little work on how exposure to these kinds of drugs may alter COMT protein expression. In one study, ten daily injections of 2 mg/kg METH significantly increased expression of both isoforms of COMT in the heart of adult male mice, both 24 h and 7 days after the final injection [55]. In the current study, however, COMT expression in the PFC was decreased 24 h after the final injection in both male and female rats following nine days of exposure to 3 mg/kg METH. This was contrary to our *a priori* hypothesis that METH would increase COMT expression in all groups, though it did increase expression in adolescent animals.

One possible explanation for the decrease in COMT expression within the PFC of adult animals is that cells where COMT is localized may be reduced following METH exposure. MB-COMT reportedly localizes with multiple pre- and post-synaptic markers and is present in relatively even distribution across neuronal cell bodies and dendrites within the rat cortex [55]. It is also expressed in glial cells [56] and appears to be more prominent in microglia than in astrocytes [57]. Reenilä et al. [58] and Helkamaa et al. [59] have reported COMT expression is increased in activated microglial cells, suggesting that changes in COMT expression may be indicative of immune response. There is evidence that the neuroimmune system continues to develop throughout adolescence, and that communication between the brain and the immune system may play in a role in a number of pathologies that tend to emerge around adolescence [60][61]. METH has been reported to induce neurotoxicity at high doses, and activation of microglia in response to METH has been reported in humans and in animals (for review, see [62]). Differences in COMT expression following METH exposure may thus be due to differences in microglia activity between adolescents and adults; however, it is important to note that most studies that demonstrate significant neuroimmune responses to METH utilize high doses (e.g. 5-10 mg/kg) in drug-naïve animals, and to date there have yet to be any published studies demonstrating if effects differ between adolescents and adults. Investigation via methods that allow for additional labeling of cell types that express COMT (e.g. immunohistochemistry) are needed to develop a better understanding of how and why the response in adult animals differs from that seen in adolescents, as the current study did not allow for such determination of mechanism.

Adolescent-specific changes in COMT expression following METH exposure may underlie some of the behavioral differences seen in both humans and animals between those exposed to amphetamines in adolescence and those exposed as adults. In rats exposed to 3 mg/kg amphetamine every other day from P27-P45 (adolescent group) or P85-P103 (adult group) who were then tested on an inhibitory control task at least 60 days after their last drug injection, our lab found that drug exposure induced impulsivity in adolescent-compared to adult-exposed males [63]. We have also found a greater deficit in working memory in male rats exposed to amphetamine during adolescence compared to those exposed as adults on a delayed-matching-to-position task [64]. Both impulse control and working memory are sensitive to PFC function, and both are sensitive to disruptions to dopaminergic signaling [65][66][67]. Lasting differences between adult and adolescent animals exposed to amphetamines may be due to imbalanced dopamine signaling, driven in part by differences in COMT expression.

The present work sought to identify, in both sexes, the normative adolescent development of COMT expression in brain regions known to undergo significant structural and functional changes during the transition from the juvenile period into adulthood. Secondarily, we aimed to investigate the potential for age-specific effects of METH exposure on COMT protein levels in the PFC. Importantly, the degree to which these effects generalize to other lower or higher doses, the pattern of drug administration, or to different ages in adulthood is unknown. We modeled non-therapeutic (i.e., “recreational”) misuse of METH in part because previous work has shown that individuals who are prescribed lower, therapeutically relevant doses of amphetamines to treat conditions such as Attention Deficit Hyperactivity Disorder (ADHD) do not have an increased likelihood of developing SUDs [68][69]. While it is difficult to compare directly between doses in rats versus humans, our selected dose of 3 mg/kg is typically considered moderate and has been shown to have age-dependent effects [47][64][70] in the absence of adverse reactions (e.g., porphyrin staining and piloerection; [71]) or neurotoxicity [72).

Additionally, the differences discussed in this report are limited to the relatively immediate timeframe following drug exposure – how, or if, these differences may persist following withdrawal is unclear, though previous work in mice has shown that COMT changes in the heart persist up to 7 days after the final dose. In the context of COMT development as a potential mechanism behind altered drug taking behaviors following adolescent exposure, lasting changes to drug self-administration following METH exposure may more closely link differences in dopamine clearance to drug use outcomes. Future work that utilizes a range of doses and measures both acute and long-term effects on COMT expression may further elucidate specific vulnerabilities to dopaminergic insults in adolescence and whether they alter adult COMT expression and/or drug taking behaviors. It would be further beneficial to measure COMT function across adolescence in a longitudinal manner, i.e. in the same animals as they develop. Future studies that utilize, for instance, microdialysis measures with and without COMT inhibitors as in [25] may provide important evidence for individual, rather than group, changes in COMT expression. Use of COMT inhibitors like tolcapone may also provide support for a mechanistic role in drug taking behaviors if utilized in a drug self-administration context.

In conclusion, METH exposure in adolescence leads to opposing results in COMT expression compared to those seen in adult exposure. The exact mechanism by which COMT expression is being modulated following METH exposure is unclear and remains in need of further examination. One possible explanation is a change in neuroimmune response to METH in these two ages, and future research into altered immune response during adolescence could help develop a clearer picture of specific changes within the PFC that lead to COMT expression shifts.

## Statement of Ethics

Experimental procedures were approved by the Institutional Animal Care and Use Committee at the University of Illinois, Urbana-Champaign, and were consistent with the Principles of Laboratory Animal Care (NIH Publication no. 85–23).

## Author Contributions

LKC was involved in all aspects of study design, execution, data analysis, and write up. JMG was involved in all aspects of study design and write up.

## Data Availability

The data that support the findings of this study are openly available at the Illinois Data Bank (https://doi.org/10.13012/B2IDB-7839940_V1).

